# Opposing roles for SNAP23 and SNAP25 in mediating MR1 trafficking and antigen presentation

**DOI:** 10.64898/2026.02.18.706493

**Authors:** Se-Jin Kim, Corinna A. Kulicke, David M. Lewinsohn, Elham Karamooz

## Abstract

MHC class I-related protein 1 (MR1) is a highly conserved antigen presenting molecule that presents small molecule metabolites derived from diverse microbial pathogens to mucosal-associated invariant T (MAIT) cells. We have shown previously that MR1 traffics through endosomal compartments via soluble N-ethylmaleimide-sensitive factor attachment protein receptor (SNARE) proteins, including Syntaxin 4 and vesicle-associated membrane protein (VAMP) 4. Here, we investigate the role of synaptosome-associated proteins (SNAPs), which pair with Syntaxins and VAMPs to form functional SNARE complexes, in MR1-mediated antigen presentation. Among SNAP homologs, we identify that SNAP23 contributes to the presentation of *Mycobacterium tuberculosis* (Mtb)-derived antigens and loss of SNAP23 reduces the number of MR1-containing vesicles during infection. In contrast, SNAP25 suppresses MR1 presentation for both intracellular pathogens Mtb and *Mycobacterium avium*, as well as extracellular pathogen *Candida albicans*. This study demonstrates opposing roles for SNAP23 and SNAP25 in MR1 antigen presentation to MAIT cells, and extends our understanding of how SNAP family proteins regulate MR1 trafficking.

## Introduction

Major histocompatibility complex (MHC) class I-related protein 1 (MR1) is a highly conserved MHC class Ib molecule that presents small molecule metabolites to mucosal-associated invariant T (MAIT) cells^1–3^. MR1 is broadly expressed in all nucleated cells and displays metabolites derived from microbial pathogens such as *Mycobacterium tuberculosis* (Mtb), as well as from host metabolism, drug molecules, and environmental sources^4,5^. MR1 presentation of some of these metabolites activates MAIT cells, promoting cytokine production to drive immune defense against infections or maintain tissue homeostasis. Previous work from our lab demonstrated that MR1 traffics through endosomal compartments to present antigens during intracellular Mtb infection^6^. More recently, pharmacologic inhibition of endosomal calcium release from two-pore channels specifically disrupted MR1-mediated presentation of Mtb while leaving HLA-Ia and exogenous antigen presentation unaffected^7^. Subsequent study identified the calcium-sensing trafficking proteins Synaptotagmin (Syt) 1 and Syt7 as potential downstream effectors of endosomal calcium release, specifically mediating MR1 presentation of Mtb by shuttling MR1 from Mtb-containing vacuoles to the cell surface for antigen presentation^8^.

Synaptosome-associated proteins (SNAPs) are key components of the soluble N-ethylmaleimide-sensitive factor attachment protein receptor (SNARE) complexes and function as essential partners of Syts in coordinating calcium-dependent membrane fusion at the plasma membrane^9–11^. Specifically, in neurons, interactions between Syt1/SNAP25 and Syt7/SNAP23 are critical for efficient vesicle exocytosis^11,12^. Among the known SNAP isoforms, SNAP23 and SNAP25 are the most extensively studied and play central roles in intracellular vesicular trafficking across diverse cell types^13^. SNAP23 is ubiquitously expressed and mediates a variety of membrane fusion events. In adipocytes, SNAP23 facilitates GLUT4 exocytosis through the interaction with Syntaxin 4^14^. In dendritic cells, it recruits the endosomal recycling complex, a rich source of MHC class I molecules, to cross-presenting phagosomes^15^. In contrast, SNAP25 is primarily expressed in neuronal cells.

While its main function in neurons is to drive regulated exocytosis, SNAP25 also participates in the endosomal recycling pathway, mediating trafficking of sorting endosomes to recycling endosomes in epithelial cells^16^. As SNAP23 and SNAP25 share significant sequence homology, emerging evidence suggests that they may compete for common binding partners and hence can modulate one another to support vesicular transport in a context-dependent manner.

In this study, we identified distinct roles for SNAP23 and SNAP25 in MR1-mediated antigen presentation in bronchial epithelial BEAS-2B cells. Among the four SNAP homologs in vertebrates, SNAP23 and SNAP25 uniquely regulate MR1 antigen presentation through opposing mechanisms. SNAP23 selectively promotes MR1 presentation of Mtb and maintains the number of MR1-containing vesicles. Furthermore, loss of SNAP23 alters MR1 cellular distribution, leading to a reduction in the total number of MR1+ vesicles at baseline and with Mtb infection. In contrast, loss of SNAP25 enhances MR1 presentation of Mtb, *Candida albicans* (*C. albicans*), and *Mycobacterium avium* (*M. avium*), which correlates with increased MR1 surface stabilization in the presence of MR1 ligand. Collectively, these findings establish SNAP23 and SNAP25 as important but functionally opposing regulators of MR1 trafficking and antigen presentation.

## Results

### SNAP homologs differentially mediate the presentation of mycobacterial antigens

In vertebrates, the SNAP family consists of SNAP23, SNAP25, SNAP29, and SNAP47. These proteins are critical for membrane fusion events such as regulated exocytosis and intracellular trafficking^13^. Therefore, we hypothesized that members of the SNAP family may contribute to antigen presentation given the role of their binding partners, Syt1 and Syt7, in MR1 antigen presentation. We first quantified the expression of *SNAP23, SNAP25, SNAP29*, and *SNAP47* in bronchial epithelial BEAS-2B cells using RT-qPCR. *SNAP23, SNAP29*, and *SNAP47* transcripts were abundantly expressed, whereas *SNAP25* transcripts were detected at comparatively lower levels (Figure 1A).

**Figure 1.**
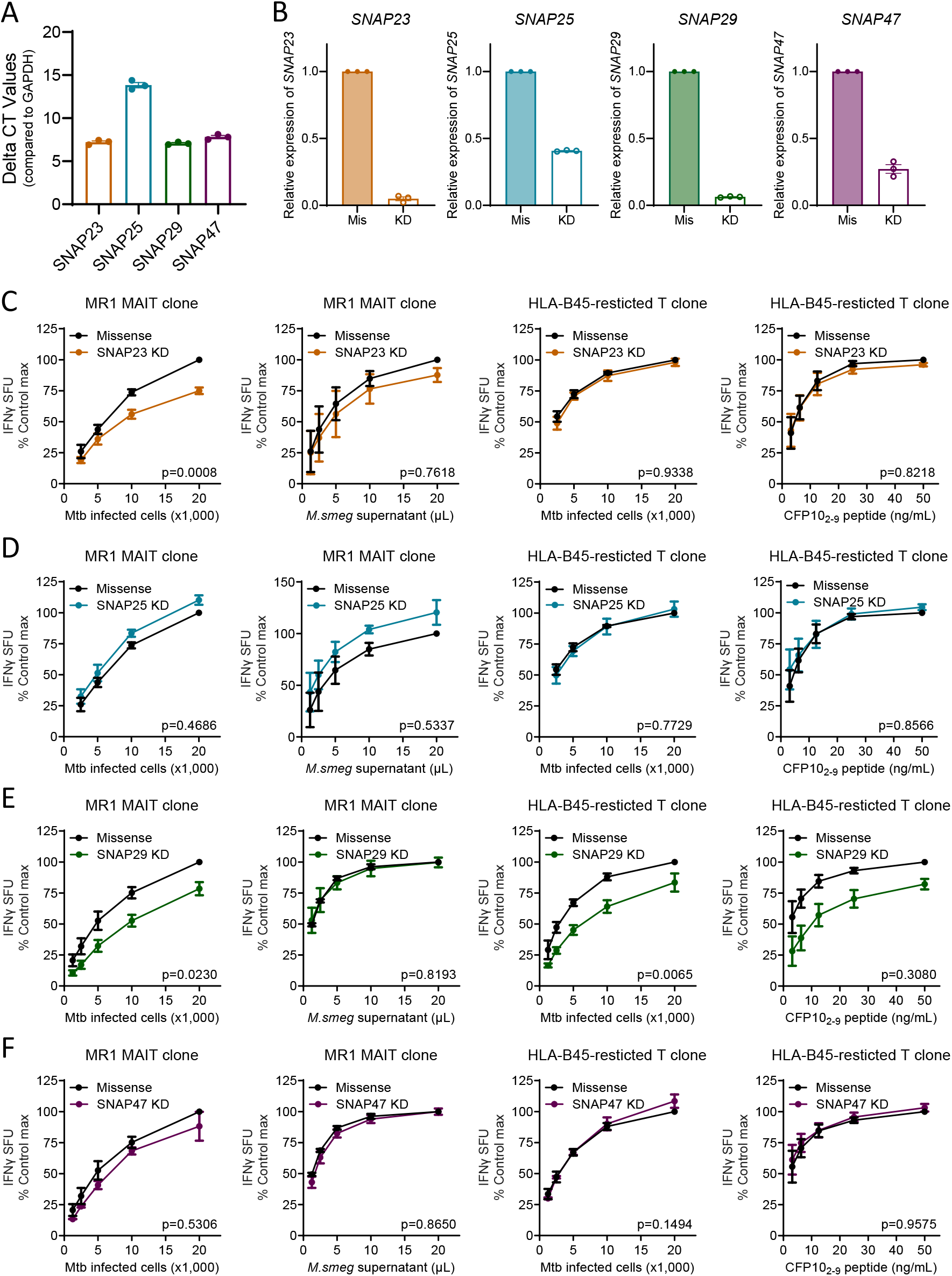
SNAP homologs differentially mediate the presentation of mycobacterial antigens. (A) Relative gene expression levels of *SNAP23, SNAP25, SNAP29*, and *SNAP47* compared to *GAPDH* in BEAS-2B cells. (B) Knockdown efficiency after 48 hours of knockdown with missense (Mis) or gene-specific (KD) siRNA for SNAP23, SNAP25, SNAP29, and SNAP47. (C-F) IFN-γ response by T cell clones (MAIT and HLA-B45-restricted) co-cultured with BEAS-2B cells following siRNA knockdown of (C) SNAP23, (D) SNAP25, (E) SNAP29, or (F) SNAP47. Cells were either infected overnight with H37Rv Mtb (MOI=8) or incubated with exogenously added antigens (*M. smeg* supernatant and CFP10_2-9_ peptide). IFN-γ response is measured by ELISpot and represented as spot forming units (SFU). All data are plotted as mean±SEM and pooled from three independent experiments. Experiments were performed in parallel for (C-D) and (E-F). For (C-F), the means of technical duplicates were pooled and normalized to Control (Missense) at the highest antigen concentration. Non-linear regression analysis comparing best-fit values of top and EC50 were used to calculate p-values by extra sum-of-squares F test.

To understand the functional contribution of each SNAP homolog to antigen presentation, we performed small-interfering RNA (siRNA)-mediated knockdown of *SNAP23, SNAP25, SNAP29*, and *SNAP47*. Knockdown efficiency was confirmed by RT-qPCR, with >90% reduction for *SNAP23* and *SNAP29*, ∼60% reduction for *SNAP25*, and ∼70% reduction for *SNAP47* transcripts (Figure 1B). Despite trying a different siRNA targeting SNAP25, the knockdown efficiency was ∼55%. Following siRNA-mediated knockdown, cells were infected with H37Rv Mtb overnight or incubated with *Mycobacterium smegmatis* (*M. smeg*) supernatant or the CFP10_2-9_ peptide, the minimal epitope for the HLA-B45-restricted T cell clone. Cells were then co-cultured with either a human MAIT or an HLA-B45-restricted T cell clone, and T cell-dependent IFN-γ release was measured (Figure 1C-F). We found that SNAP23 knockdown reduced MR1 presentation of Mtb, without affecting MR1 presentation of *M. smeg* supernatant or HLA-B45 presentation (Figure 1C). SNAP25 knockdown showed a trend toward enhanced MR1 presentation for both Mtb and *M. smeg* supernatant, although these changes did not reach statistical significance (Figure 1D). Interestingly, SNAP29 knockdown impaired presentation of Mtb to both MAIT and HLA-B45-restricted T cell clones, indicating an effect on both MR1 and HLA-Ia presentation (Figure 1E). SNAP47 knockdown had no impact on MR1 or HLA-B45 antigen presentation (Figure 1F). These findings suggest that SNAP homologs differentially regulate antigen presentation, with SNAP23 and SNAP25 having specific but opposing effects on MR1 antigen presentation.

### Opposing roles of SNAP23 and SNAP25 in MR1-dependent antigen presentation of intracellular and extracellular pathogens

To further define the role of SNAP23 and SNAP25 in MR1-dependent antigen presentation, we used the CRISPR/Cas9 system to knock out SNAP23 and SNAP25 in BEAS-2B cells. We confirmed gene knockout (KO) efficiency in clonal cell lines with Inference of CRISPR Edits (ICE) analysis and measured changes in mRNA transcript levels by RT-qPCR^17^. ICE analysis showed high editing efficiency of 90% for SNAP23 and 100% for SNAP25 (Figure S1A). Transcript levels of SNAP23 and SNAP25 were greatly diminished in the KO cell lines (Figure S1B). No SNAP23 protein was detected in the KO cells (Figure S1C). SNAP25 protein expression could not be reliably detected due to nonspecific binding by two commercially available antibodies. Consistent with siRNA-mediated knockdown results, SNAP23 KO cells significantly reduced MR1 presentation of Mtb with no changes in the presentation of *M. smeg* supernatant (Figure 2A, orange). Compared to siRNA-mediated SNAP25 knockdown, SNAP25 KO cells resulted in a significant increase in MR1 presentation of both Mtb and *M. smeg* supernatant (Figure 2A, teal). Neither SNAP23 nor SNAP25 KO cells altered HLA-B45-dependent antigen presentation (Figure S1D).

**Figure 2.**
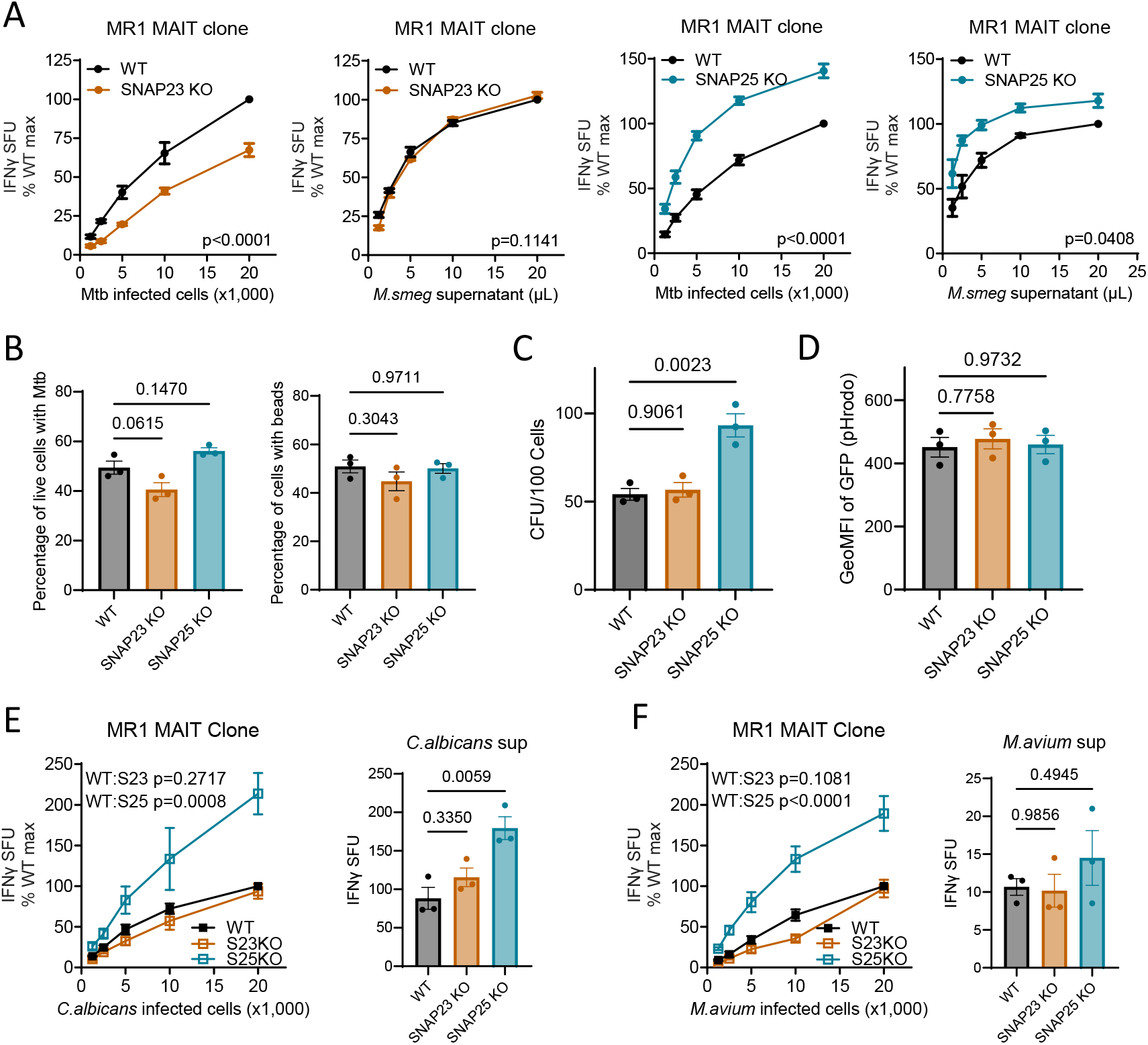
Opposing roles of SNAP23 and SNAP25 in MR1-dependent antigen presentation of intracellular and extracellular pathogens. Clonal SNAP23 KO and SNAP25 KO cells lines were generated from BEAS-2B cells (Figure S1). (A) Cells were either infected overnight with H37Rv Mtb (MOI=8) or incubated with *M. smeg* supernatant. IFN-γ response is measured by ELISpot and represented as spot forming units (SFU). The means of technical duplicates were pooled from three independent experiments and normalized to WT at the highest antigen concentration. Non-linear regression analysis comparing best-fit values of top and EC50 were used to calculate p-values by extra sum-of-squares F test. (B) Percent of cells with Mtb (left) or beads (right) that are GFP+ after infection with mEmeraldRFP-AuxMtb (MOI=8) or incubation with yellow-green fluorescent beads (ratio=8) overnight. (C) Colony forming units (CFU) of H37Rv Mtb (MOI=8) in cells after overnight infection. (D) GeoMFI of GFP of cells treated with pHrodo dextran green. (E-F) Cells were either infected with microbes or incubated with supernatants of *C. albicans* and *M. avium*. Cells were infected with *C. albicans* (MOI=0.2) for 90 minutes or with *M. avium* (MOI=22) for overnight. 50 µL of supernatants were added for both *C. albicans* and *M. avium*. IFN-γ response is measured by ELISpot and represented as spot forming units (SFU). The means of technical duplicates were pooled from three independent experiments, and non-linear regression analysis comparing best-fit values of top and EC50 were used to calculate p-values by extra sum-of-squares F test. For (B-F), ordinary one-way ANOVA with Dunnett’s multiple comparisons test were used to analyze significant differences. All data are plotted as mean±SEM.

Since SNAP23 has been implicated in phagosome formation in macrophages^18^, we next investigated whether SNAP23 or SNAP25 contributes to Mtb uptake or phagosome formation in BEAS-2B cells. We conducted Mtb uptake assays using an auxotrophic strain of Mtb (AuxMtb), which constitutively expresses a green fluorescent protein^19,20^. We observed no significant differences in the uptake of AuxMtb for both SNAP23 and SNAP25 KO cells compared to wildtype cells (Figure 2B, left). Similarly, we also incubated the cells with inert yellow-green fluorescent beads overnight to distinguish from Mtb-driven intracellular responses. We observed no changes in bead uptake (Figure 2B, right). Although SNAP23 and SNAP25 did not impact the uptake of AuxMtb and beads, we sought to assess the viability of intracellular Mtb. We performed colony-forming unit (CFU) assays in lysed cells after overnight infection with H37Rv Mtb^7^. Notably, there was a significant increase in Mtb viability in SNAP25 KO cells compared to wildtype control (Figure 2C). To determine whether the increased Mtb viability was due to changes in endosomal pH, we measured organelle acidity using pHrodo dextran green and detected no significant differences (Figure 2D). These findings suggest that SNAP23 and SNAP25 are not involved in the uptake of Mtb or beads, but SNAP25 potentially plays a role in the viability of Mtb, impacting the number of available Mtb ligands for presentation, through mechanisms independent of endosomal acidification.

To determine whether the role of SNAP23 and SNAP25 in MR1 presentation is generalizable to other microbes, we next tested *C. albicans* and *M. avium* as an extracellular and intracellular pathogen, respectively. SNAP23 KO showed no effect on MR1 presentation of *C. albicans* and its supernatant. In contrast, SNAP25 KO augmented MR1 presentation under both conditions (Figure 2E). Similarly, we infected SNAP23 and SNAP25 KO cells with *M. avium* overnight, and found enhanced MR1 presentation of *M. avium* in SNAP25 KO cells only (Figure 2F). *M. avium* supernatant was very weakly antigenic, eliciting only 8-20 compared to 100-200 IFN-γ spots, and showed no differences in presentation for both SNAP23 and SNAP25 KO cells. Altogether, these data suggest opposing roles of SNAP23 and SNAP25 in MR1-dependent antigen presentation. SNAP23 selectively promotes MR1 presentation of Mtb, while SNAP25 negatively regulates MR1 presentation across both intracellular and extracellular microbes.

### SNAP25 suppresses 6-FP-dependent MR1 surface stabilization

Given that SNAP25 KO increased MR1-mediated antigen presentation, we next assessed whether this effect is associated with an increase in MR1 surface stabilization in the presence of a ligand. We transfected SNAP25 KO cells with tetracycline-inducible GFP-tagged MR1 (tetMR1-GFP) and induced MR1-GFP expression with doxycycline before treating the cells with either NaOH (solvent control) or 6-formylpterin (6-FP) to promote MR1 translocation from the endoplasmic reticulum (ER) to the cell surface. We found that as in WT BEAS-2B cells, the total MR1 (surface and intracellular) expression as measured by GFP signal intensity did not change in SNAP25 KO cells irrespective of 6-FP (Figure 3A). In contrast, SNAP25 KO cells displayed enhanced MR1 surface translocation in response to 6-FP, while there were no changes in HLA-Ia surface levels (Figure 3B-C). This suggests that SNAP25 may be involved in regulating stability of surface MR1.

**Figure 3.**
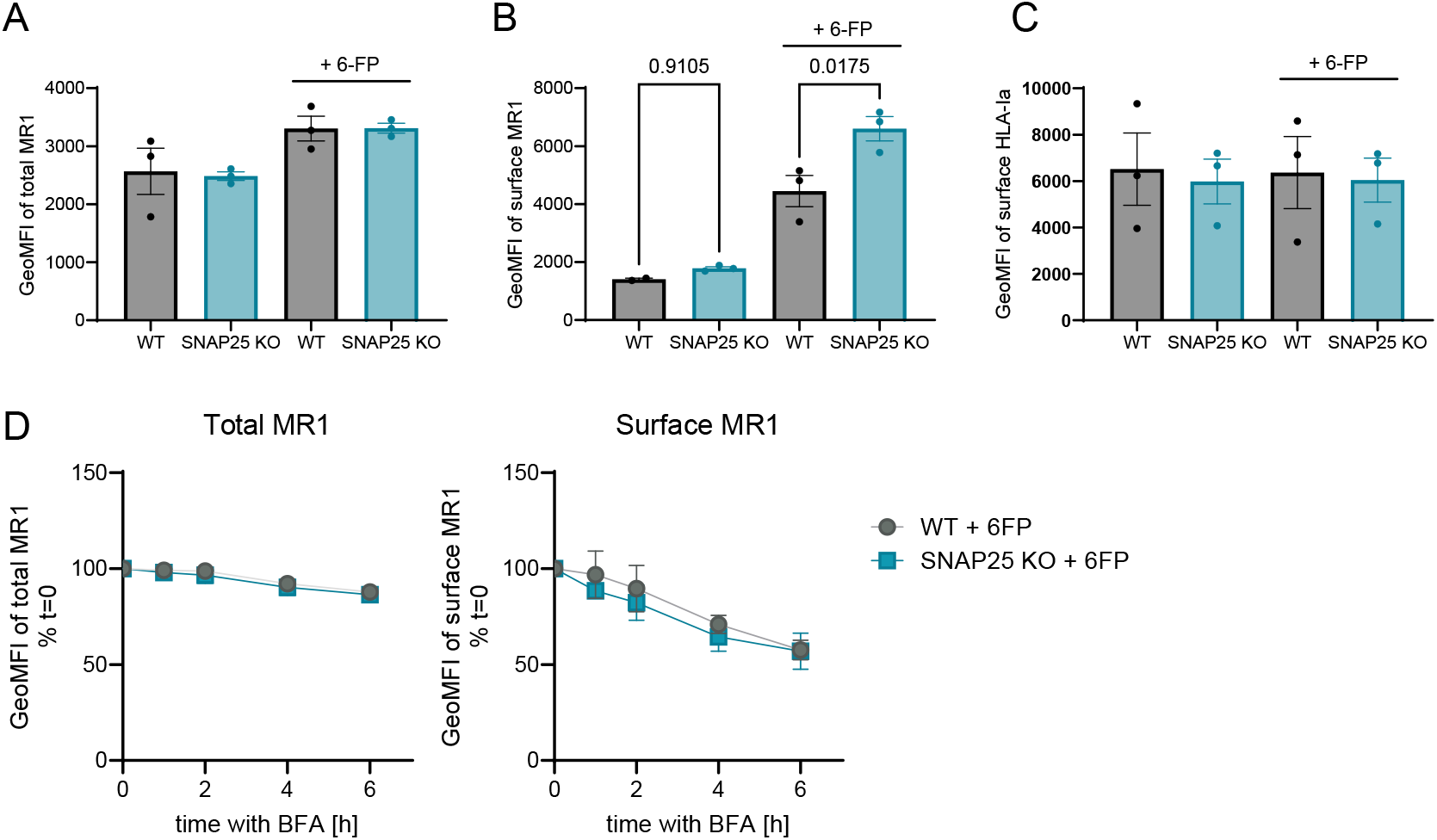
SNAP25 suppresses 6-FP-dependent MR1 surface stabilization. WT and SNAP25 KO BEAS-2B cells were transfected with TETMR1-GFP and incubated with doxycycline overnight. 6-FP or NaOH (solvent control) was added the next day for at least 18 hours. (A-C) Cells were analyzed by flow cytometry. Representative of three independent experiments with pooled geometric mean fluorescence intensity (GeoMFI) of GFP+ cells on total MR1 (A), surface MR1 (B) and surface HLA-Ia expression (C). For BFA decay assays, cells were incubated with brefeldin A (BFA) for indicated time periods after induction of cells with doxycycline and 6-FP. (D) GeoMFI of GFP+ cells on total MR1 (left) and surface MR1 expression (right) were measured by flow cytometry. Representative of four independent experiments with pooled GeoMFI for each time point. Non-linear regression analysis on one-phase exponential decay curves and comparing one curve fit to both data sets by extra sum-of squares F test. All data are plotted as mean±SEM.

To further determine whether SNAP25 affects presentation of newly synthesized MR1 or internalization of MR1 from the cell surface, we performed a Brefeldin A (BFA) assay to block anterograde transport of new MR1 and measured the decay of surface MR1^21^. Total MR1 levels measured by GFP expression slightly decreased for both WT and SNAP25 KO cells after BFA treatment. Surface MR1 levels decreased to ∼50% over the course of 6 hours, but there were no significant differences between WT and SNAP25 KO cells (Figure 3D). These findings indicate that SNAP25 does not alter MR1 internalization, but instead increases the rate at which loaded MR1 reaches the cell surface.

### Quantity of MR1 vesicles depends on SNAP23

In contrast to SNAP25, which suppresses MR1 presentation of both intracellular and extracellular microbes, SNAP23 specifically promotes MR1 presentation of Mtb (Figure 2). To understand the role of SNAP23 in MR1 trafficking and cellular distribution, we generated a clonal SNAP23 KO cell line from BEAS-2B MR1 KO cells stably transduced with a doxycycline-inducible MR1-GFP plasmid (BEAS-2B:TET-MR1GFP)^21^. RT-qPCR, ICE analysis, and Western blot verified complete loss of *SNAP23* transcript and protein (Figure S2A-C). SNAP23 KO cells exhibited a marked reduction in both MR1- and HLA-B45 presentation of Mtb (Figure 4A). However, the magnitude of effect was greater in the presentation of Mtb to the MAIT cell clone than HLA-B45-restricted T cell clone. We next assessed whether SNAP23 KO had impaired MR1 surface stabilization in the presence of a ligand. Flow cytometry showed no differences in MR1 surface translocation or HLA-Ia expression between WT and SNAP23 KO after 6-FP treatment (Figure 4B). These results suggest that SNAP23 does not mediate the translocation of 6-FP loaded MR1 to the plasma membrane.

**Figure 4.**
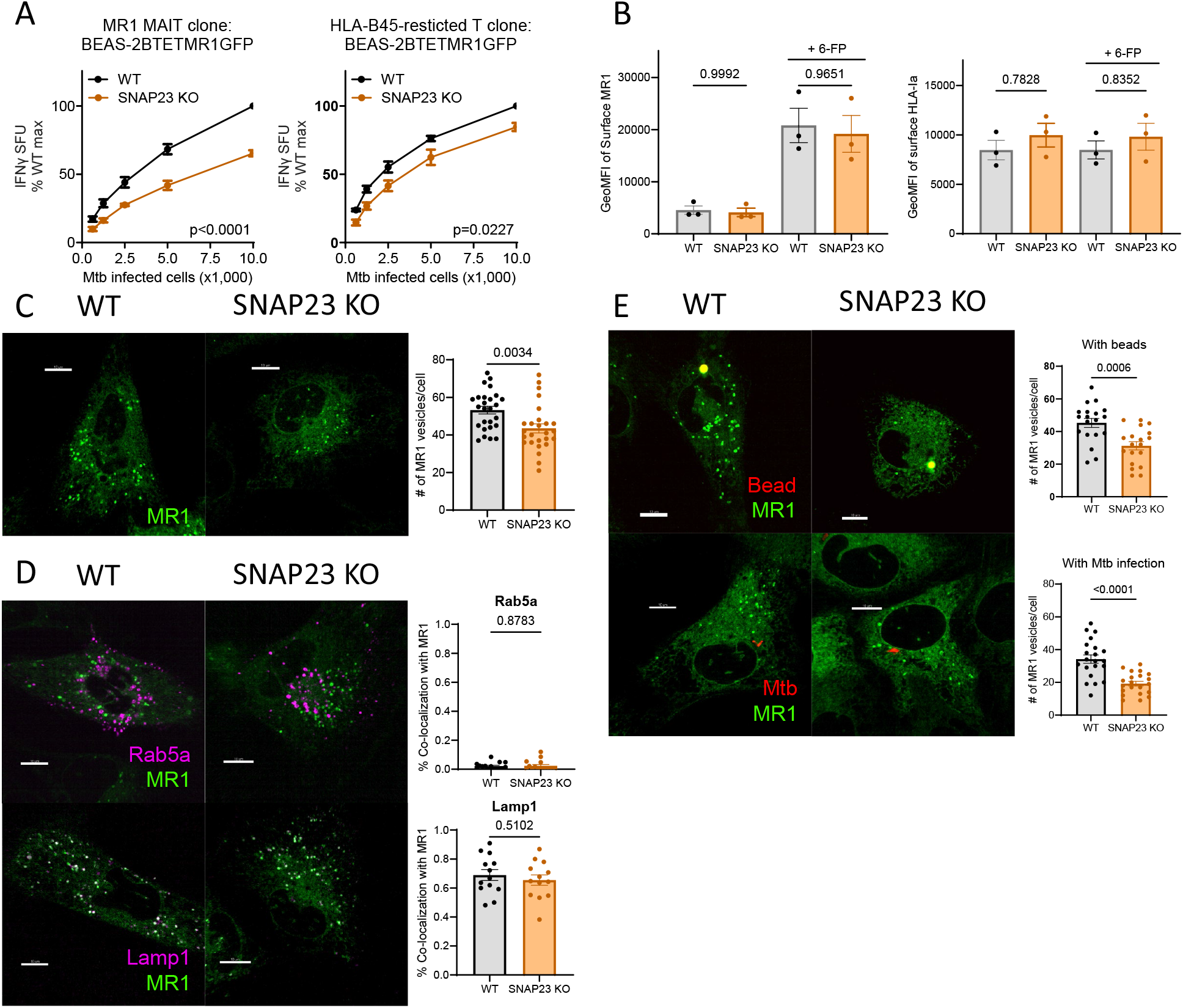
Quantity of MR1 vesicles depends on SNAP23. A clonal SNAP23 KO cell line was generated in the background of BEAS-2B MR1KO:tetMR1-GFP clone D4 (WT) cells (Figure S2). (A) Cells were infected overnight with H37Rv Mtb (MOI=8). IFN-γ response by T cell clones (MR1- and HLA-B45-restricted) is measured by ELISpot and represented as spot forming units (SFU). The means of technical duplicates were pooled and normalized to WT at the highest antigen concentration. Non-linear regression analysis comparing best-fit values of top and EC50 were used to calculate p-values by extra sum-of-squares F test. For MR1 surface expression, cells were incubated with doxycycline overnight. 6-FP or NaOH (solvent control) was added the next day for at least 18 hours. (B) Cells were then analyzed for geometric mean fluorescence intensity (GeoMFI) of GFP+ cells on surface MR1 and surface HLA-Ia expression. (C-E) Representative images of WT and SNAP23 KO BEAS-2B MR1KO:tetMR1-GFP cells at (C) steady state, (D) infected overnight with CellLight Rab5a (top) or LAMP1 (bottom), (E) incubated overnight with beads (top) or infected with AuxMtb (bottom). (C, E) Total number of MR1 vesicles in each cell. (D) Percent co-localization of Rab5a and LAMP1 with MR1 vesicles. Each dot represents one cell. p-values were analyzed by two-tailed unpaired Student’s t-test. All data points are pooled and representative of three independent experiments, and plotted as mean±SEM. All scale bars represent 10 µm.

To investigate whether SNAP23 facilitates MR1 vesicular trafficking and cellular distribution, we performed fluorescence microscopy on SNAP23 KO BEAS-2B:TET-MR1GFP cells. Interestingly, SNAP23 KO cells displayed a significant reduction in the number of MR1+ vesicles per cell at baseline (Figure 4C). To determine whether this reduction altered MR1 trafficking, we infected cells with baculoviruses expressing RFP-tagged Rab5a (early endosomes) or LAMP1 (lysosomes). Co-localization analysis showed that MR1 vesicles in both WT and SNAP23 KO cells localized predominantly with LAMP1 (∼70%) and minimally with Rab5a (<5%), indicating that loss of SNAP23 does not reroute MR1 vesicles to alternative compartments (Figure 4D). Finally, we examined whether the number of MR1+ vesicles changes in response to phagocytosis. Both bead uptake and intracellular Mtb infection also reduced the number of MR1+ vesicles in SNAP23 KO cells (Figure 4E). Together, these data suggest that SNAP23 is critical for maintaining quantity of MR1+ vesicles, which may contribute in efficient MR1-mediated presentation of Mtb.

## Discussion

SNARE-mediated membrane fusion is crucial for intracellular vesicular trafficking, facilitating the transport of cargo, including antigen presenting molecules, between organelles and to the plasma membrane. Syts are calcium-sensing trafficking proteins that interact with SNARE complexes to regulate vesicle exocytosis^10,22^. Specific protein pairings, such as Syt1/SNAP25 and Syt7/SNAP23, are important for efficient neurotransmitter release^11,12^. Beyond their role in neuronal exocytosis, recent studies show that Syts also mediate the translocation of antigen presenting molecules, such as MHC class II and MR1, to the plasma membrane^8,23^. Becker *et al*. demonstrated that Syt7 drives translocation of MHC class II from late endosomes to the plasma membrane in dendritic cells, while our group showed that Syt1 and Syt7 specifically mediate MR1 presentation of Mtb-derived antigens^8,23^. In this study, we extend these observations by identifying distinct and opposing roles of the Syt partners SNAP23 and SNAP25 in MR1-mediated antigen presentation and highlight potential mechanisms through which they regulate MR1 trafficking pathways.

Although SNAP23 and SNAP25 share 55% amino acid sequence homology^13^, our findings show that they play opposing roles in MR1 antigen presentation in bronchial epithelial cells. SNAP23 specifically promotes MR1 presentation of Mtb, whereas SNAP25 inhibits MR1 presentation of both intracellular and extracellular microbes. While both proteins are known for their functional role in promoting exocytosis, prior studies in other cell types also report opposite roles^24–26^. Kunii *et al*. identified that the loss of SNAP23 decreases granule fusion in exocrine cells, but increases fusion in endocrine cells^24^. This finding suggests that the function of SNAP proteins can differ depending on the cell type and may explain the inhibitory role of SNAP25 we observe in MR1 presentation in bronchial epithelial cells. Moreover, other studies show that SNAP23 and SNAP25 differ in membrane affinity, may compete for shared binding partners, and modulate one another’s function^25–27^. Salaün *et al*. demonstrated that SNAP23 contains an additional cysteine residue within its membrane-targeting domain compared to SNAP25, enhancing its association with the plasma membrane^27^. In addition, SNAP23 can inhibit formation of SNARE complexes by preventing SNAP25 from binding Syntaxin1A, and depletion of SNAP23 enhances SNAP25 binding to voltage-gated calcium channels and increases exocytosis^25,26^. These studies suggest that the loss of either SNAP homolog can alter formation of SNARE complexes and membrane fusion events, potentially influencing MR1 trafficking and presentation as well.

We have previously shown that MR1 trafficking and presentation are mediated by endosomal proteins and found differences in the presentation of intracellular Mtb and exogenous antigen such as *M. smeg* supernatant^6,28^. For example, Syntaxin 4 does not affect MR1 presentation of Mtb, but it contributes to the presentation of exogenous antigens^28,29^. Recent work further demonstrates that endosomal calcium signaling and calcium-sensitive trafficking proteins are important in mediating MR1 presentation of Mtb^7,8^. Pharmacologic inhibition of calcium release using small molecule *trans*-Ned-19 specifically reduces MR1 antigen presentation of Mtb and *M. avium*^7^. Our findings also show that SNAP23 specifically regulates MR1 presentation of Mtb. To determine whether these pathways and function of trafficking proteins are shared for other microbes, we examined the role of SNAP23 and SNAP25 in MR1 presentation of extracellular pathogen *C. albicans* and intracellular pathogen *M. avium*. Unexpectedly, SNAP23 KO cells did not alter MR1 presentation of either pathogen or their supernatants, whereas SNAP25 KO cells augmented MR1 presentation of all sources of antigen except for *M. avium* supernatant, which was weakly antigenic. These findings indicate that SNAP23 and SNAP25 do not contribute to MR1 presentation uniformly across intracellular and extracellular microbes, nor through endosomal calcium signaling. Instead, their roles appear to depend on the specific pathogen, which has unique antigens and endosomal proteins that mediate MR1 presentation.

Mtb mainly resides in membrane-bound phagosomes, but the precise location of MR1 antigen processing and loading remains unknown^30–32^. Prior work demonstrated that SNAP23 regulates phagosome formation and maturation^18^. Although our findings showed that SNAP23 KO cells in bronchial epithelial cells did not alter Mtb uptake or phagocytosis of beads, we observed a significant decrease in the number of MR1+ vesicles at baseline. The progression of these MR1+ vesicles through the endosomal pathway remained unchanged, as loss of SNAP23 did not affect their co-localization with Rab5a or LAMP1. Furthermore, addition of beads or infection with Mtb reduced the number of MR1+ vesicles, suggesting that these vesicles may serve as intermediates in the presentation of intracellular Mtb antigens. This concept is further supported by our recent work showing that MR1+ vesicles, whether loaded or unloaded, potentially traffic from Mtb-containing vacuoles to the cell surface for antigen presentation^8^. Follow-up studies that characterize antigen-loading status of MR1 and time-course experiments tracking these MR1+ vesicles during intracellular Mtb infection will be critical for understanding how SNAP23 supports MR1-mediated antigen presentation.

In this study, we show that SNAP23 and SNAP25 have opposing roles in MR1 antigen presentation. SNAP23 is important for maintaining the quantity of MR1 vesicles, while SNAP25 suppresses MR1 antigen presentation, as demonstrated by SNAP25 KO cells showing augmented MR1 antigen presentation of extracellular and intracellular pathogens as well as exogenously delivered antigens. We also observed increased Mtb viability, which may enhance antigen availability and thereby promote presentation. However, there were no changes in organelle acidity or properties of the cells that might explain this phenotype, underscoring the need for additional mechanistic studies. Defining their shared binding partners as well as contributions to constitutive versus regulated exocytosis may clarify how SNAP23 and SNAP25 have opposing roles. Overall, our findings reveal that SNAP23 and SNAP25 differentially regulate MR1 trafficking, defining a new role for the SNAP family in antigen presentation.

## Methods

### Bacterial Strains and Cell Lines

H37Rv *Mycobacterium tuberculosis* (Mtb) and *Mycobacterium avium* Chester (both ATCC) were grown in Middlebrook 7H9 Broth supplemented with Middlebrook ADC (BD), 0.05% Tween-80 (OmniPur), and 0.5% glycerol (Thermo Fisher Scientific). *Candida albicans* Berkhout (ATCC) was grown in Brain Heart Infusion (BHI) Broth (Sigma). All microbes for infection were used from frozen glycerol stocks after passaging 20 times through a tuberculin syringe (BD) with a 27-gauge needle before infection. Multiplicity of Infection (MOI) of 8 (Mtb), 22 (*Mycobacterium avium* Chester), and 0.2 (*Candida albicans*) were determined by titering frozen stocks and used for IFN-γ ELISpot assays. Supernatants were obtained by culturing *M. avium* in supplemented 7H9 Broth and *C. albicans* in BHI Broth for 24 hours and passing the supernatant through a 0.22 μm filter. Supernatants were used from frozen aliquots stored at -80°C and also incubated in medium to confirm that they did not have viable bacteria. ΔleuD ΔpanCD double auxotroph Mtb (AuxMtb), an attenuated strain of H37Rv Mtb, was gifted by Dr. William Jacobs. A modification of AuxMtb was generated to constitutively express the green fluorescent protein mEmerald and the tetracycline (tet)-inducible red fluorescent protein (mEmeraldRFP-AuxMtb)^19,20^. For AuxMtb, frozen glycerol stock was thawed, grown for several days in 7H9 broth with 0.05% Tween-80 and leucine/pantothenate, and used for infection based on OD_600_ readings. MOIs of 5 (for fluorescence microscopy) and 8 (for Mtb uptake assay) were used. *Mycobacterium smegmatis* (*M. smeg*, mc^2^155 strain) supernatant was obtained by culturing *M. smeg* for 24 hours in 7H9 broth, passing the supernatant through a 0.22 μm filter, and concentrating it using a 10kDa Amicon filter (Millipore Sigma). *M. smeg* supernatant was used from frozen aliquots stored at -80°C.

BEAS-2B cells were obtained from ATCC. BEAS-2B MR1KO:tetMR1-GFP clone D4 was used to generate SNAP23 KO BEAS-2B MR1KO:tetMR1-GFP cells^21^. BEAS-2B cells were cultured in DMEM (Gibco) supplemented with 10% heat-inactivated fetal bovine serum (GeminiBio) and L-Glutamine (Gibco).

### Ribonucleoprotein-mediated gene knockouts

All knockouts were generated in BEAS-2B and BEAS-2B MR1KO:tetMR1-GFP clone D4 cells using a CRISPR Gene Knockout Kit (Synthego) per the manufacturer’s protocol on CRISPR editing of immortalized cell lines (Synthego). Knockouts and monoclonal cell lines were generated as described before^8^.

### Human Subjects

All samples were collected with informed consent, and all experiments were conducted according to protocols approved by the Institutional Review Board at Oregon Health & Science University (IRB00000186). Peripheral blood mononuclear cells (PBMCs) were obtained by apheresis from healthy adults and used to expand T cell clones as described below. Human serum was also obtained from healthy adults.

### T cell clones

T cell clones were rapidly expanded by co-culturing with irradiated allogenic PBMCs and allogenic lymphoblastoid cell lines in RPMI1640 medium (Gibco) containing 10% human serum and anti-CD3 (30 ng/mL, clone OKT3, eBioscience). Recombinant IL-2 (2 ng/mL, OHSU Inpatient Pharmacy) was added the following day and every 2-3 days thereafter. On Day 5, T cell clones were washed to remove anti-CD3 and frozen down at 1-1.5e6 cells/mL after at least 11 days. New freezeback stocks were validated before use by comparing IFN-γ response to a previous freezeback. Two T cell clones were previously characterized and used throughout the study: a MAIT cell clone (D426-G11) and an HLA-B45-restricted T cell clone (D466-A10)^33,34^.

### Reagents and chemicals

Doxycycline (Sigma-Aldrich) was resuspended to 2 mg/mL in H_2_O and used at 2 µg/mL unless specified. 6-FP (Schirck’s laboratories) was resuspended in 0.01 M NaOH at 1 mg/mL and used at 100 µM. Phytohemagglutinin (PHA, Roche) was resuspended at 10 mg/mL in RPMI1640 medium (Gibco) supplemented with 10% human serum, 2% L-glutamine, and 0.1% gentamicin for ELISpot assays. 16% Paraformaldehyde (PFA, Electron Microscopy Sciences) was diluted in PBS to appropriate concentrations. pHrodo Dextran Green (Thermofisher) was used at 33 µg/mL. CFP10_2-9_ peptide was obtained from Genemed Synthesis and resuspended in DMSO at 5 mg/mL.

### ELISpot assays

IFN-γ enzyme-linked immunosorbent spot (ELISpot) assays were conducted to determine T cell dependent IFN-γ responses to microbes and antigens as previously described^8^. For Mtb and *M. avium* infection, 3e4 or 4e4 BEAS-2B cells were infected with H37Rv Mtb (MOI=8) and *M. avium* (MOI=22) overnight. For *C. albicans* infection, 3e4 BEAS-2B cells were infected with MOI of 0.2 for 90 minutes, washed twice, and incubated with media containing Antibiotic Antimycotic Solution (Sigma-Aldrich) overnight. Infected BEAS-2B cells were serially diluted and plated in duplicate. For supernatants and peptides, 1e4 BEAS-2B cells were plated in duplicate. Cells were incubated with *M. smeg* supernatant, *C. albicans* supernatant, *M. avium* supernatant, or CFP10_2-9_ peptide as indicated for each experiment for 1 hour at 37°C and 5% CO_2_. For all experiments, 1e4 T cell clones (MAIT cell clone (D426-G11) or HLA-B45-restricted T cell clone (D466-A10)) were added and co-cultured for at least 18 hours at 37°C and 5% CO_2_. PHA (Roche) was used as a positive control. All cells and reagents were resuspended in the blocking buffer (RPMI (Gibco) containing 10% human serum, 2% L-glutamine, and 0.1% gentamycin (Gibco)) for plating. IFN-γ ELISpots were developed and enumerated as previously described^8^.

### CFU assays

Cells infected with H37Rv Mtb (MOI=8) overnight were lysed in ultrapure water the next day. Serial dilutions of 1:10 and 1:100 with PBS + 0.05% Tween-80 were plated in triplicates on 7H10 agar supplemented with glycerol and Middlebrook ADC. Plates were incubated at 37°C and 5% CO_2_ for 12 days. Number of colonies were counted to determine colony-forming units per 100 cells.

### siRNA knockdown

Small-interfering RNAs targeting SNAP23 (s16709), SNAP25 (s13189), SNAP29 (s17861), SNAP47 (s42061), and a missense control (4390844) were obtained from Thermo Fisher Scientific. 1.5e5 BEAS-2B cells were plated in 6-well plates (Corning) and transfected the next day with 50 nM siRNA using Lipofectamine RNAiMAX (Invitrogen) at 80% confluency. Cells were used for ELISpot assays or RNA extraction at 48 hours post-transfection.

### RNA isolation, cDNA synthesis, and qPCR analysis

Total RNA was isolated using the RNeasy Plus Mini Kit (Qiagen) and reverse transcribed into cDNA using the High-Capacity RNA-to-cDNA Kit (Applied Biosystems) according to the manufacturer’s instructions. Quantitative RT-qPCR was performed on a Step One Plus Real-Time PCR System (Applied Biosystems) using TaqMan Universal PCR Master Mix (Life Technologies). Taqman FAM-MGB probes for *SNAP23* (Hs01047496_m1, Hs01047498_m1), *SNAP25* (Hs00938957_m1, Hs00938966_m1), *SNAP29* (Hs00191150_m1), and *SNAP47* (Hs00369605_m1) were obtained from ThermoFisher Scientific. Samples were run in triplicates and gene expression levels were normalized to *GAPDH* (Hs02758991_g1) of the same sample. Relative expression of the sample was compared to missense control.

### Western blot

Equal cell numbers were lysed in lysis buffer containing 0.5% NP-40 (Chromotek) and protease inhibitor cocktail (Roche) for 1 hour on ice. Whole cell lysates were collected after centrifugation and combined with loading buffer (Invitrogen) and reducing agent (Invitrogen). Samples were loaded on a 4-20% Mini-Protean TGX gel (Bio-Rad). The gel was run at 120V for 1 hour and transferred to a polyvinylidene fluoride membrane (Millipore) at 100V for 1 hour in the cold room. The membrane was blocked with Odyssey blocking buffer (Li-Cor) for 1 hour at room temperature. The membrane was stained with primary antibody against SNAP23 (1:1000; Abcam, ab131242, rabbit) and loading control vinculin (1:1000; Bio-Rad, MCA465GA, mouse) in Odyssey antibody diluent (Li-Cor) at 4°C overnight. The membrane was washed in PBS with 0.1% Tween-20 (Affymetrics) and incubated with secondary antibodies against mouse (1:10,000; Li-Cor, 925-68072) and rabbit (1:10,000; Li-Cor, 926-32213) conjugated to different fluorophores. The membrane was imaged on an Odyssey CLx imager (LI-COR).

### Flow cytometry assays

Mtb uptake assays were conducted as previously described^8^. Cells were infected with mEmeraldRFP-AuxMtb (MOI=8) overnight, stained with Live/Dead Fixable Dead Cell Stain Kit (Thermo Fisher), and fixed in 4% PFA before analysis. Similarly, for bead uptake assay, 1e5 BEAS-2B cells were plated in a 12 well plate and incubated at 37°C and 5% CO_2_ for at least 3 hours. 1 μm carboxylate-modified polystyrene yellow-green fluorescence latex beads were diluted in PBS to 4.75e7 beads/mL. Latex beads were incubated with BEAS-2B cells at a ratio of 8. Cells were washed with PBS and fixed in 1% PFA. For MR1 surface expression assays, SNAP23 or SNAP25 KO cells were generated in the background of WT BEAS-2B and subsequently transfected with TET-MR1GFP^35^ or SNAP23 or SNAP25 KO cells were generated from BEAS-2B MR1KO:tetMR1-GFP clone D4. These cells were plated in a 6-well plate and treated with 2 μg/mL doxycycline overnight at 37°C and 5% CO_2_. 100 μM 6-FP or 0.01 M NaOH (solvent control) was added the next day for at least 18 hours. Cells were stained with Live/Dead Fixable Dead Cell Stain Kit, APC-conjugated anti-HLA Ia antibody (clone W6/32; BioLegend, 311410), APC-conjugated anti-MR1 antibody (clone 26.5; BioLegend, 361108), and APC-conjugated isotype control antibody (clone MOPC-173; BioLegend, 400222) in a FACS buffer (PBS containing 2% human serum, 2% goat serum, and 0.5% FBS). Cells were washed and fixed in 2% PFA. The BFA decay assay was done as previously described^21^. Cells in the background of BEAS-2B transfected with TET-MR1GFP or BEAS-2B MR1KO:tetMR1-GFP clone D4 were treated with 2 or 4 μg/mL doxycycline, respectively, overnight at 37°C and 5% CO_2_. The next day, 100 μM 6-FP or 0.01 M NaOH (solvent control) was added along with fresh doxycycline. Medium containing doxycycline and 6-FP was washed and replaced with new medium containing doxycycline and 10 μg/mL BFA at indicated time points. Cells were stained for MR1 surface expression as described above. All data was obtained with LSR II (BD) cytometer at the OHSU Flow Cytometry Shared Resource and analyzed with FlowJo software version 10 (TreeStar).

### Fluorescence Microscopy

BEAS-2B cells were plated on either 4-well or 8-well #1.5 glass bottom chamber slides (Nunc) and incubated with 2 μg/mL doxycycline at 37°C and 5% CO_2_. After at least 3 hours of incubation, cells were incubated with latex beads (Sigma-Aldrich), CellLight BacMam 2.0 (Invitrogen), or infected with mEmeraldRFP-AuxMtb overnight. 2 μm carboxylate-modified polystyrene red fluorescence latex beads were diluted in PBS to 4.75e6 beads/mL. Latex beads were incubated with BEAS-2B cells at 1:1 ratio. For determining MR1 co-localization with endosomal compartments, CellLight for Rab5a-RFP (C10587) or LAMP1-RFP (C10597) was added. For Mtb infection, cells were infected with mEmeraldRFP-AuxMtb (MOI=5) in the presence of leucine and pantothenate. Cells were imaged the next day in an unbiased manner based on MR1-GFP expression. All images were acquired on a motorized Nikon TiE stand with a Yokogawa W1 spinning disk unit and a high-powered Agilent laser-emission filter (405nm-445/50nm, 488nm-525/36nm, 561nm617/73nm) at the OHSU Advanced Light Microscopy Core. A 100x (NA 1.49) objective was used and images were captured with an Andor Zyla 5.5 sCMOS camera with 2by2 camera binning.

### Image Analysis

All images were processed and analyzed on Imaris 7 (Bitplane). MR1 vesicles were enumerated with the “Spots” function by defining the region of interest for each cell, estimated XY diameter as 1 μm, and quality of spots greater than 20. Co-localizations were analyzed using the “Spots” function and “spots colocalization” MatLab Xtension module as previously described^6^.

### Statistical Analysis

All data were analyzed in Prism 10 (GraphPad). At least three independent experiments were performed and plotted as mean±SEM. For ELISpot assays, mean data from technical replicates were pooled. Non-linear regression analysis was conducted with no constraints to compare differences in best-fit values of top and EC50 between the curves. p-values were calculated using agonist versus response (three parameters) and extra sum-of-squares F test. Unless otherwise indicated, two tailed unpaired t-test (for two groups), a one-way ANOVA with Dunnett’s multiple comparisons test (for three groups), or a one-way ANOVA with Tukey’s multiple comparisons test (for four groups) were conducted for statistical analysis. A p value of <0.05 was considered statistically significant. For BFA decay assays, best fit parameters for one-phase exponential decay curves were determined by least-squares regression. The fit of one curve fit to both data sets was compared with the fit of individual curves fit to each data set using the extra sum-of-squares F test.

## Acknowledgments

The research reported in this publication used computational infrastructure supported by the Office of Research Infrastructure Programs, Office of the Director, of the National Institutes of Health under Award Number S10OD034224. The content is solely the responsibility of the authors and does not necessarily represent the official views of the National Institutes of Health. We acknowledge the assistance of the Oregon Clinical & Translational Research Institute, which is supported by the National Center for Advancing Translational Sciences, National Institutes of Health, through Grant Award Number UL1TR002369. We acknowledge expert technical assistance by staff in the Advanced Light Microscopy Core (RRID:SCR_009961) in the Department of Neurology and Jungers Center, as well as OHSU Flow Cytometry and Monoclonal Antibody Shared Resource Core Facility (RRID:SCR_009974) at Oregon Health and Science University. We thank staff at the Vollum DNA Sequencing Core. The contents do not represent the views of the U.S. Department of Veterans Affairs or the United States Government. Lastly, we are grateful to Dr. William Jacobs for sharing the Mtb auxotroph and Dr. Shogo Soma for generation of the mEmeraldRFP-AuxMtb.

## Author Contributions

The experiments presented were conceptualized by SK, CK, EK, and DML. SK performed experiments and analyzed data. DML and EK supervised the work. CK, EK, and DML provided advice and technical expertise. All authors contributed to revising and reviewing the manuscript. All authors approved the final version of the manuscript.

---

**Supplementary Figure 1. Characterization of SNAP23 KO and SNAP25 KO in the background of BEAS-2B cells**

(A) Genome editing efficiency and percent distribution of individual indels of SNAP23 and SNAP25 KO BEAS-2B cells as determined by Sanger sequencing and ICE analysis^17^.

(B) *SNAP23* (left) and *SNAP25* (right) transcript levels compared to *GAPDH* measured by RT-qPCR.

(C) SNAP23 protein expression measured by western blot for WT and clonal SNAP23 KO BEAS-2B cell line.

(D) Cells were either infected overnight with H37Rv Mtb (MOI=8) or incubated with CFP10_2-9_ peptide and co-cultured with HLA-B45-restricted T cell clones. IFN-γ response is measured by ELISpot and represented as spot forming units (SFU). All data are plotted as mean±SEM and pooled from three independent experiments. The means of technical duplicates were pooled and normalized to WT at the highest antigen concentration. Non-linear regression analysis comparing best-fit values of top and EC50 were used to calculate p-values by extra sum-of-squares F test. Experiments were performed in parallel with those shown in Figure 2A.

**Supplementary Figure 2. Validation of SNAP23 KO in the background of BEAS-2B MR1KO:tetMR1-GFP cells**

(A) *SNAP23* transcript levels compared to *GAPDH* measured by RT-qPCR.

(B) Genome editing efficiency and percent distribution of individual indels of SNAP23 KO BEAS-2B MR1KO:tetMR1-GFP cells as determined by Sanger sequencing and ICE analysis^17^.

(C) SNAP23 protein expression measured by western blot for WT and SNAP23 KO BEAS-2BMR1KO:tetMR1-GFP cells.

