## Supplementary figures and images for "Opposing roles for SNAP23 and SNAP25 in mediating MR1 trafficking and antigen presentation"

### Supplemental Figure 1 and 2

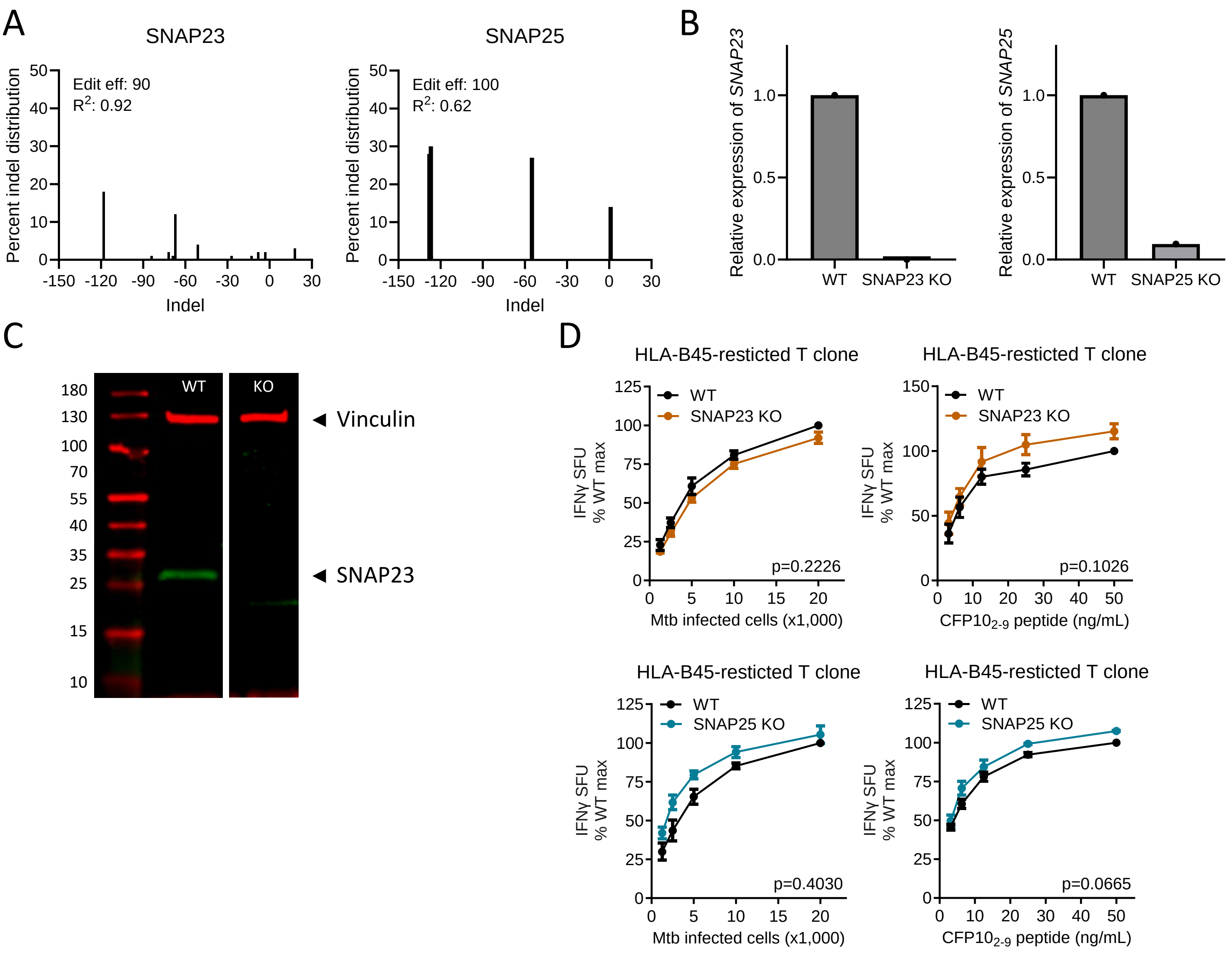

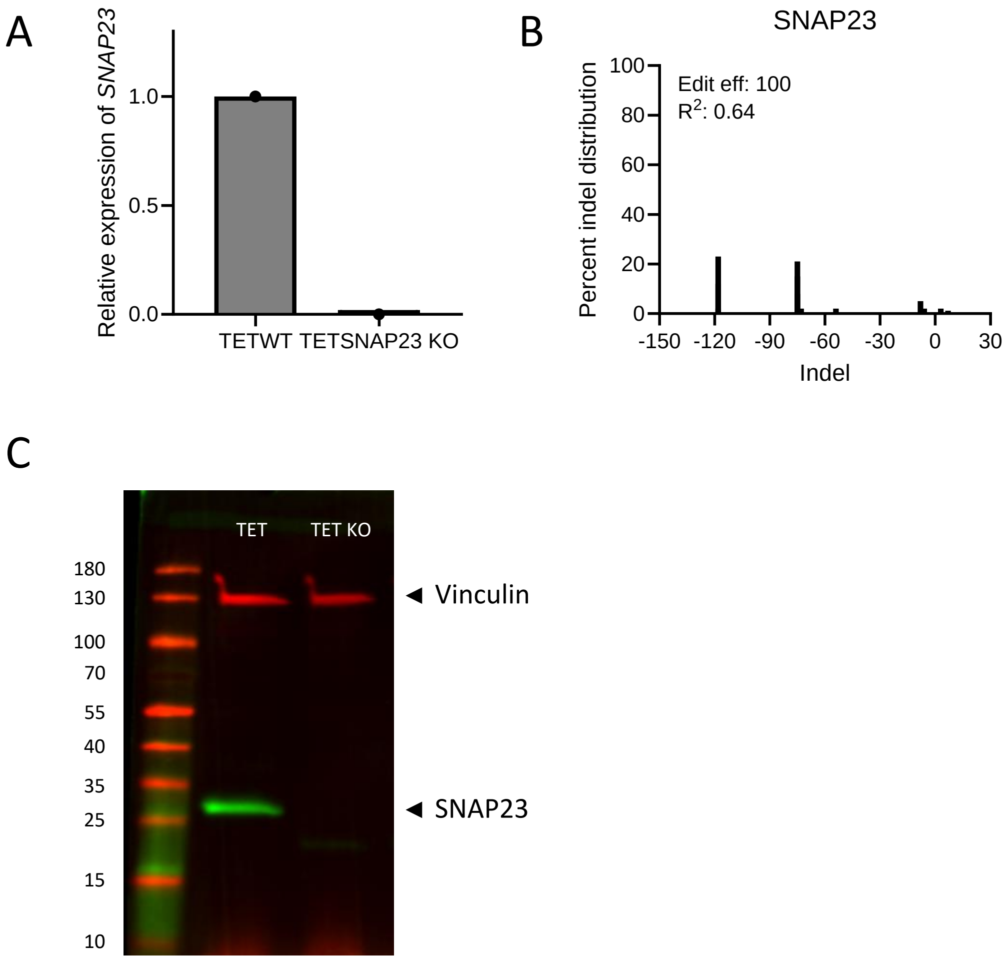
